# Development and evaluation of a pan-fungal lateral flow device for the rapid identification of pathogen class in microbial keratitis

**DOI:** 10.64898/2026.04.29.721399

**Authors:** Leonie Fingerhut, Sheelagh Duncan, Bethany Mills

## Abstract

**Purpose:** To develop a pan-fungal lateral flow device (LFD); evaluating device performance with samples obtained from *ex vivo* porcine cornea infection models.

**Methods:** Fungal β-glucan (PF1), human CLEC7A/Dectin-1/CLECSF12 protein (Fc Tag; PF2), and fungal melanin (PF3) antibodies were evaluated for binding to clinically relevant fungal and bacterial species (*Aspergillus flavus, Fusarium keratoplasticum, Candida albicans, Pseudomonas aeruginosa, Staphylococcus aureus*) by immunofluorescence staining. PF1 and PF2 were evaluated in proof-of-concept, in-house LFD strips using cultured pathogens and *ex vivo* porcine corneal infection samples. The lead antibody (PF1) was validated in a commercially-developed prototype LFD.

**Results:** PF1 and PF2 discriminated target fungi from bacteria by immunofluorescence microscopy and in-house LFD strips. The lead candidate PF1 demonstrated good sensitivity (0.75) and specificity (0.94) with cultured fungal hyphae. Samples obtained from infected *ex vivo* cornea by clinically relevant methods confirmed excellent sensitivity (scrapes: 1.00, swabs: 0.94) and specificity (scrapes: 1.00, swabs: 0.83). The commercially-developed PF1-LFD prototype achieved perfect sensitivity (1.00) and specificity (1.00) when detecting and discriminating fungi from non-fungal *ex vivo* corneal swab samples.

**Conclusions:** Feasibility of pan-fungal LFD application in microbial keratitis diagnosis was demonstrated – using β-glucan as a pan-fungal target and a clinically relevant microbial keratitis *ex vivo* model. LFDs were able to differentiate fungal from bacterial samples, detect antigen present in corneal swabs, and provide a read-out within 20 minutes. Sensitivity and specificity values are comparable to currently used diagnostic tests.

**Translational Relevance:** Early discrimination of fungal keratitis cases is important to adapt treatment, and improve patient outcome.

## Introduction

Microbial keratitis (MK) is a common infection of the cornea following an initial microinjury of the ocular surface. It is caused by a broad spectrum of pathogens, exceeding 680 bacterial, fungal, viral and amoeboid species.^1^ The disease affects over two million people annually,^2^ leaving many with long-term visual impairment, making it the second leading cause of unilateral blindness after cataracts.^2,3^

The predominant pathogen class causing MK varies depending on geographic location, with bacterial cases being most frequent in high-income countries across Europe and North America.^4,5^ In contrast, fungal cases dominate in tropical regions.^6,7^ The geographic variation is caused by climate and contact lens use, but also abundancy of workers exposed to fungal spores, such as people working in agriculture.^3^ Globally, fungal keratitis cases are estimated to exceed one million per year,^8^ and a trend in increasing prevalence is reported.^7^ Fungal keratitis is associated with worse outcomes compared to bacterial infections, as well as prolonged treatment times and healthcare expenses.^9-12^ Sixty percent of fungal keratitis patients remain with visual impairment, and 50% of UK patients require surgical intervention.^10^ It is therefore an acute ophthalmic emergency. Delayed diagnosis and initiation of appropriate treatments is a leading risk factor for a poor outcome.^9^ Yet, regions with the highest incidence (rural areas of low- and middle-income countries) have the least access to tertiary care facilities with diagnostic laboratories. ^7,8^

Diagnostic laboratories are crucial in fungal keratitis diagnosis as clinical presentation is similar to other pathogen classes.^13^ In a recent study, the importance of diagnosis was highlighted as 76.9% of fungal keratitis cases were treated with the wrong antimicrobial when assumptions of the causative pathogen were made from clinical presentation.^14^ It is essential to diagnose fungal cases in early stages to adjust treatment and improve clinical outcomes. Microbial culture and smear microscopy are the current gold-standard,^7,15^ however, culture may take 7-14 days for fungal diagnosis and suffers poor sensitivity.^4,5,16^ Smear microscopy provides same-day results, with a sensitivity of 61-94% and specificity of 91–97% for detection of fungi,^11^ but requires invasive corneal scrapes, microscopy equipment, and skilled healthcare professionals for execution. Molecular technologies are very sensitive, which can lead to false-positive results from commensal conjunctival contaminants, but most importantly their expense prohibits their wide application.^7,17^ Despite wide access to diagnostic methods in the UK, the average interval between presentation and confirmation of fungal aetiology was 31 days.^18^ Crucially, ∼50% of common diagnostic testing fails to identify the causal pathogen. ^4,5,16^

Therefore, novel diagnostic methods are urgently needed to provide easy and rapid identification of the pathogen class outside of tertiary care facilities. Recently, the first reported feasibility study demonstrating the use of repurposed commercially available lateral flow devices (LFDs) for the identification of *Aspergillus* causing MK was performed.^19^ Results were available within 30 minutes of sample collection, and a sensitivity/specificity of 89% and 95% was achieved. As this test is limited to the detection of *Aspergillus* cases, we now aim to identify fungal cases in general, using a pan-fungal LFD approach – which becomes the most important deciding factor when initiating antimicrobial treatments for MK. The majority of fungal keratitis cases are caused by filamentous fungi *Aspergillus* spp. and *Fusarium* spp. or yeast *Candida* spp.,^6,10^ hence we aim to detect all of these in one test.

Recently, the Fungitell (1,3)-β-D-glucan assay was demonstrated for this application,^20^ however this assay requires laboratory personnel and equipment, limiting its use to tertiary care. Major advantages of a LFD as a diagnostic tool include its simple use, rapid diagnosis, low sample volume, applicability at point-of-care, long shelf life, stability at room temperature, and the low cost.^21^ Applicability of LFDs for easy, rapid, and widely used diagnosis has been demonstrated for instance during the COVID-19 pandemic, or in pregnancy testing. Moreover, several fungal lateral flow tests were recently developed for the rapid detection of *Mucorales*,^22^ *Rhizopus arrhizus*,^23^ or *Coccidioides*.^24^ However, to our knowledge, there are no pan-fungal LFDs reported to date.

## Material and Methods

### Pathogen culture

The fungal strains *Aspergillus flavus* ATCC200026 and *Fusarium keratoplasticum* ATCC36031 were sourced from ATCC. Spores were harvested after 3-5 days growth at 30°C on potato dextrose (PD, Millipore 70139) agar growth plates, using phosphate-buffered saline (PBS) with 0.01% Tween 80 (Sigma P1754). Spore concentration was determined by haemocytometer counting and adjusted to 10^5^ spores mL^-1^. Hyphae were grown from freshly harvested spores by static incubation at 30°C in PD broth (Millipore P6685) overnight.

The fungal strain *Candida albicans* strain ATCCMYA2876 was sourced from ATCC. *C. albicans* was prepared from a single colony picked from a PD agar growth plate and inoculated into PD broth to grow overnight at 37°C under constant motion. The overnight culture was adjusted to OD_595_ 0.1 and further incubated until mid-log phase (OD_595_ 0.4-0.8). Yeast cells were washed three times with 0.9% sterile saline (Baxter), and concentrations were then readjusted to a final concentration of 10^5^ conidia mL^-1^ by haemocytometer counting. Conidia were stimulated to hyphal transformation by static incubation at 30°C in RPMI (Gibco 11835-063) overnight.

*S. aureus* strain IHMA2190153 was sourced from IHMA bacterial repository, *Pseudomonas aeruginosa* PAO1 was sourced from ATCC. Single colonies from Luria-Bertani (LB) agar plates were inoculated into Tryptic Soy Broth (TSB, Oxoid CM0129) in the case of *S. aureus* and into LB broth (Sigma L3147) for *P. aeruginosa*. These were grown overnight at 37°C under motion. The overnight culture was adjusted to OD_595_ 0.1 and further incubated until mid-log phase (OD_595_ 0.4-0.8). Bacteria were washed three times with 0.9% sterile saline.

### Immunofluorescence staining and microscopy

Spores of filamentous fungi (300 µL of *A. flavus* at 7.5×10^3^ spores mL^-1^ and *F. keratoplasticum* at 3×10^4^ spores mL^-1^) were seeded into poly-D-lysine (Sigma P6407) coated µ-slide 8 well imaging chambers (Ibidi 80821) in PD broth and incubated overnight at 30°C to allow hyphae growth. *C. albicans* yeast, derived from a fresh subculture grown in PD broth, was washed three times and resuspended in RPMI to an OD_595_ 0.05. 200 µL of the yeast suspension was added per well into µ-slide imaging chambers and incubated overnight at 30°C. Bacterial strains (*P. aeruginosa* and *S. aureus*) were grown overnight in LB or TSB broth respectively, subcultured, washed, and added to the imaging chambers at an OD_595_ of 0.1. All samples were fixed using 4% paraformaldehyde (Thermo Scientific J19943-K2).

Prior to immunofluorescence staining, samples were blocked with 1% bovine serum albumin (BSA, Sigma A4503) for 30 minutes at room temperature. As primary pan-fungal antibodies, **PF1:** rabbit fungal β-glucan monoclonal antibody (Abexxa abx023999, final concentration 2 µg/mL); **PF2:** recombinant human CLEC7A/Dectin-1/CLECSF12 protein (Fc Tag) (Sino Biological 10215-H01H, final concentration 1 µg/mL); or **PF3:** mouse anti-fungal melanin recombinant IgG (Creative Biolabs FAMAB-0160WJ, final concentration 10 µg/mL), were applied in blocking buffer for one hour. Isotype controls included rabbit IgG (R&D systems AB-105-C, final concentration 2 µg/mL), mouse IgG2a from murine myeloma (Merck M5409), and a no primary antibody control for the human dectin-1 Fc protein. Then, samples were carefully washed three times, and secondary antibody (either Alexa Fluor^TM^ 488 goat anti-mouse IgG (H+L) (A11001, final concentration 4 µg/mL), or Alexa Fluor® 488 goat anti-human IgG (H+L) (Mol. Probes A11013, final concentration 4 µg/mL), or Alexa Fluor® 488 F(ab’)_2_ fragment of goat anti-rabbit IgG (H+L) (Invitrogen A11070, final concentration 4 µg/mL)) in PBS was added for one hour at room temperature in the dark. Further washes followed, and samples were kept in PBS (bacteria and *C. albicans*) for imaging or imaged with minimal liquid to prevent fungal filaments from floating (*A. flavus* and *F. keratoplasticum*).

Microscopy was performed on a Leica SP8 confocal laser scanning microscope (HC PL APO CS2, 10x and 63x objective). Two wells were imaged per condition, and repeated independently thrice. Images were processed using ImageJ software (version v1.54k). Maximum intensity projections of z-stacks shown for *A. flavus* and *F. keratoplasticum*.

### In-house lateral flow strip assembly (s-PF1-LFD and s-PF2-LFD)

A Universal Lateral Flow Assay Kit (Abcam ab270537) was used to create a custom sandwich lateral flow assay following the manufacturer’s instructions with slight modifications. In short, pan-fungal candidates **PF1** and **PF2** were conjugated to Lightning-Link® Ulfa-Tag and 40 nm Gold at concentrations of 1 mg/mL and 0.3 mg/mL, respectively. Antibody was up-concentrated using Antibody Concentration and Clean-up Kit (Abcam, ab102778) if necessary. These conjugate stocks were then further diluted in running buffer (10x Universal Running buffer diluted to 1x in dH_2_O with 0.1% BSA final) to 150 µg/mL for Lightning-Link® Ulfa-Tag conjugate and 3:10 in case of the gold conjugate. Per sample, 5 µL each of the diluted Lightning-Link® Ulfa-Tag and Gold conjugates were mixed with Gold-Biotin (1:10 diluted in running buffer) and 75 µL analyte (described below) in running buffer. These were incubated for five minutes at room temperature, before 80 µL was loaded into wells of a low-binding 96 well plate and lateral flow assay strips inserted. The liquid was soaked up the sample pad and assays (**s-PF1-LFD** and **s-PF2-LFD**) were read after 20 minutes. For each sample, an image was taken using a smartphone, as well as two independent blinded visual scores (using a standard intensity chart ranging 0-8). The images were analysed using a previously developed ratiometric method.^19^ In short, the ImageJ Gel Analyser Tool was used to determine the intensity profiles of control and test bands, from which their peak areas were quantified and the ratio of test (T) to control (C) line calculated. Using T/C values over T value alone removes ambiguity of variances of the in-house test intensities.

### Prototype LFD development

Prototype lateral flow devices (**p-PF1-LFD**) were manufactured by Lateral Dx Ltd. (Alloa, Scotland, UK). Devices were assembled using a CN140 nitrocellulose membrane (25□mm width) laminated onto a backing card and fitted with an Ahlstrom 222 top pad (27□mm), a polyester conjugate pad (15□mm), and an Ahlstrom 1281 sample pad (15□mm). The membranes were striped with **PF1** at 0.5 mg/mL, together with a goat anti rabbit IgG control line at 0.3□mg/mL, both prepared in borate buffer. **PF1** was biotinylated and subsequently conjugated to an anti-biotin gold colloid prior to application.

100 µL of sample in running buffer (PBS-Tween) was applied per lateral flow test, tests were developed for 20 minutes and then read using a commercial cube reader (LateralDx Ltd.), as per manufactures instructions.

### Preparation of samples for LFD assessment

#### Preparation of standards

Zymosan A from *Saccharomyces cerevisiae* (Sigma Z4250) was diluted in sterile Dulbecco’s PBS (dPBS; Gibco 14190-094) at a stock concentration of 10 mg/mL. The mixture was heated at 95°C for ten minutes and intermittently vortexed, and subsequently broken up by repeated aspiration through a 23G (BD Microlance^TM^ 300800) and then a 27G needle (BD Microlance^TM^ 300635). This zymosan stock was stored at 4°C for up to two weeks prior to use. On the day of measurement, the stock solution was vortexed and working solutions between 10 and 500 µg/mL were prepared in running buffer.

β-D-glucan from barley (Sigma, G6513) was diluted in dH_2_O at a stock concentration of 20 mg/mL, and then further diluted in running buffer to concentrations of 0-5 ng/mL.

#### Preparation of cultured pathogens

Fungal hyphae samples were grown as described above. The supernatant was taken off carefully, hyphae were washed once using saline, then resuspended in running buffer. The hyphae were lifted from the well bottom using a pipette tip and then homogenised to break up clumps in Precellys Evolution at 6500 rpm, two cycles at 20 seconds (Precellys P000912-LYSK1-A.0). Bacterial samples were obtained as described above and resuspended in running buffer to an OD_595_ of 0.1.

75 µL of prepared sample was used for the lateral flow assays. Supernatants were serially diluted and plated on PD or LB agar for quantification of colony forming units from pathogen samples after one (bacteria and *Candida*) or two (filamentous fungi) days incubation at 37°C (bacteria) or 30°C (filamentous fungi and *Candida*), 5% CO_2_. For CFUs of filamentous fungi, samples were serially diluted at a 1:10 ratio, plated into a 24 well plate, and growth at the highest dilution used for quantification, as these fungi do not grow in colonies. Each condition was run in duplicates and in three independent repeats.

#### Preparation of e*x vivo* porcine microbial keratitis model samples

Fresh porcine eyes were received from “pre-tank” pigs within 12 h of their slaughter from a local abattoir (Browns Food Group (Quality Pork Processors Ltd.)) and kept on ice during transportation. The extraocular tissue was removed, and the eye balls immersed in sterile phosphate-buffered saline (PBS). The eyes were further cleaned using PBS containing 2% penicillin/streptomycin (Gibco 15140122) and 2% amphotericin B (Sigma A2942) for two minutes, and then 3% povidone iodine (Merck 1.09099.1000) in PBS for two minutes. The cleaned globes were washed a further three times in antibiotic/antifungal-free PBS and maintained at 4°C overnight in PBS. Next the cornea was soaked in 20% ethanol for two minutes, then the epithelium was removed mechanically from the cornea with a scalpel blade dipped in 70% ethanol. The cornea-scleral rims were dissected from the globe, and the lens, ciliary body, iris, and trabecular meshwork were removed. The cornea-scleral rims were washed in PBS, and cleaned again using 2% penicillin/streptomycin/amphotericin B for two minutes, and then 3% povidone iodine for two minutes. Next, the cornea-scleral rims were filled with 2% [w/v] ultra-pure agarose (Invitrogen 16550-100) dissolved in PBS. Each cornea was placed epithelial side up into a well of a 6-well plate. Ten µL of fungal spores or yeast (10^5^ mL^-1^ in PBS), bacteria (10^6^ CFU mL^-1^) or sterile PBS as control, was pipetted onto the centre of the cornea. The well housing the cornea-scleral rim was filled with PBS without reaching the level of the cornea. Eyes were incubated at 30°C for 48 hours. Uninfected cornea had PBS containing antimicrobials (2% penicillin/streptomycin/amphotericin B) added into the agarose, and were incubated at 4°C to prevent microbe growth.

After incubation, swab and scrape samples were obtained from the models. Micro swabs (Puritan 25-800 R50) were dipped in running buffer, then samples were collected by swabbing over the ocular surface. The swab tip was placed immediately into 150 µL running buffer. For scrapes, the surface of the cornea was scratched three times with a 21G needle (BD Microlance^TM^ 304432) and the needle tip placed immediately into 150 µL running buffer. Each sample was vortexed for two minutes before measurements.

Lateral flow signal and CFUs were measured as above. Each condition was run in duplicates and in three independent repeats.

#### Statistical analysis

Data were analysed using GraphPad Prism 9.5.1 (GraphPad Software, San Diego, CA, USA). Statistical dependencies were determined via simple linear regression. Diagnostic performance (sensitivity, specificity, positive and negative predictive values, accuracy) were calculated with MedCalc Version 23.5.1 (MedCalc Software Ltd). Independent experiments were performed at least three times, measuring duplicates.

## Results

### LFD prototype development pipeline

To develop a pan-fungal LFD, potential antibodies and Fc-fusion proteins were first screened against clinically relevant MK pathogens by immunofluorescence microscopy. Next, antibodies successful in binding to fungi but not bacteria were then conjugated and evaluated in in-house developed, strip-based LFDs. Finally, the lead antibody was integrated into a commercially-developed pan-fungal LFD prototype for further evaluation (**Figure 1**).

**Figure 1.**
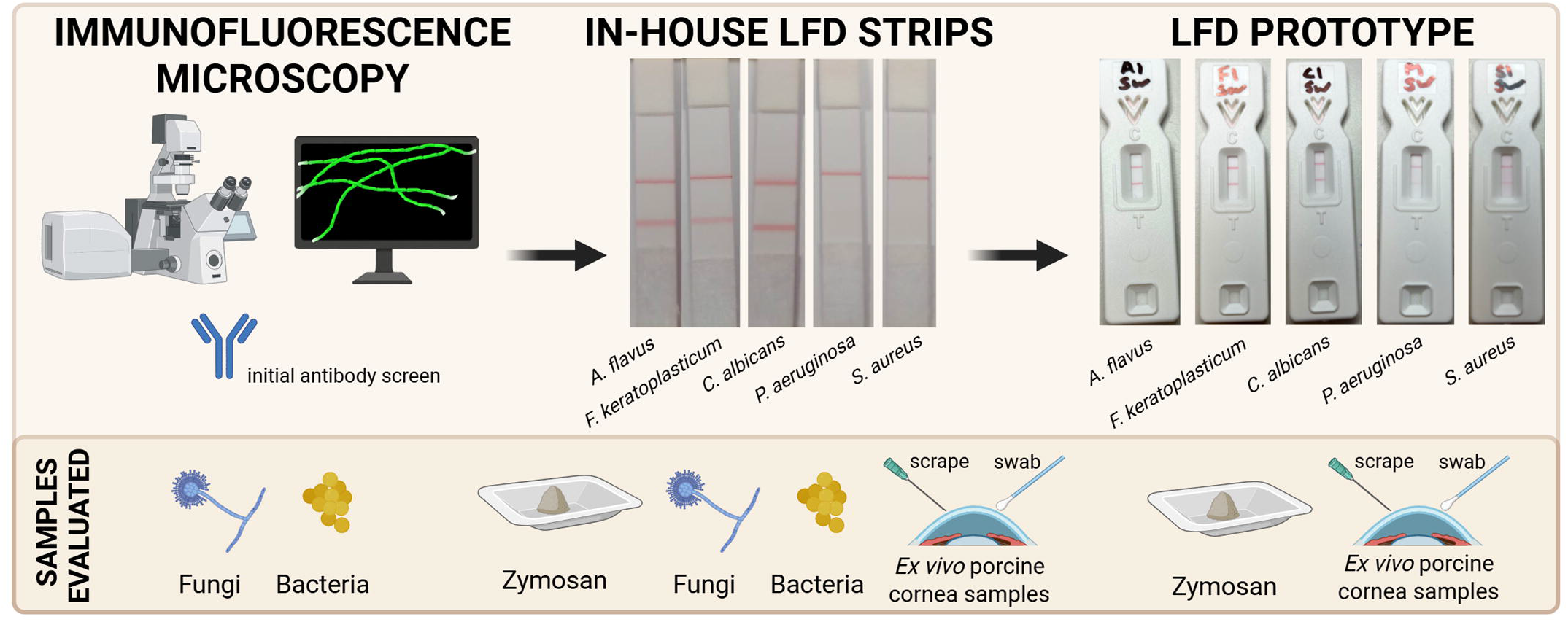
Development pipeline for pan-fungal lateral flow device (LFD) - from antibody screening by immunofluorescence microscopy, to feasibility with LFD strips developed in-house, to development of an optimised commercially-developed LFD prototype.

### Specific staining of fungal but not bacterial MK pathogens by PF1 and PF2

Commercially available antibodies and Fc-fusion proteins to common and abundant fungal cell wall components (β-glucan (**PF1, PF2**) and fungal melanin (**PF3**)) were screened against target fungi and off-target bacteria by immunofluorescence staining and microscopy. Filamentous fungi *Aspergillus flavus* and *Fusarium keratoplasticum*, as well as yeast (*Candida albicans*), resulted in positive staining with **PF1** and **PF2** (**Figure 2**). No off-target staining of *Pseudomonas aeruginosa* (gram-negative bacteria) or gram-positive *Staphylococcus aureus* (gram-positive bacteria) was detected. **PF3** did not result in specific staining of fungal hyphae. Therefore, only **PF1** and **PF2** were progressed to in-house LFD development.

**Figure 2.**
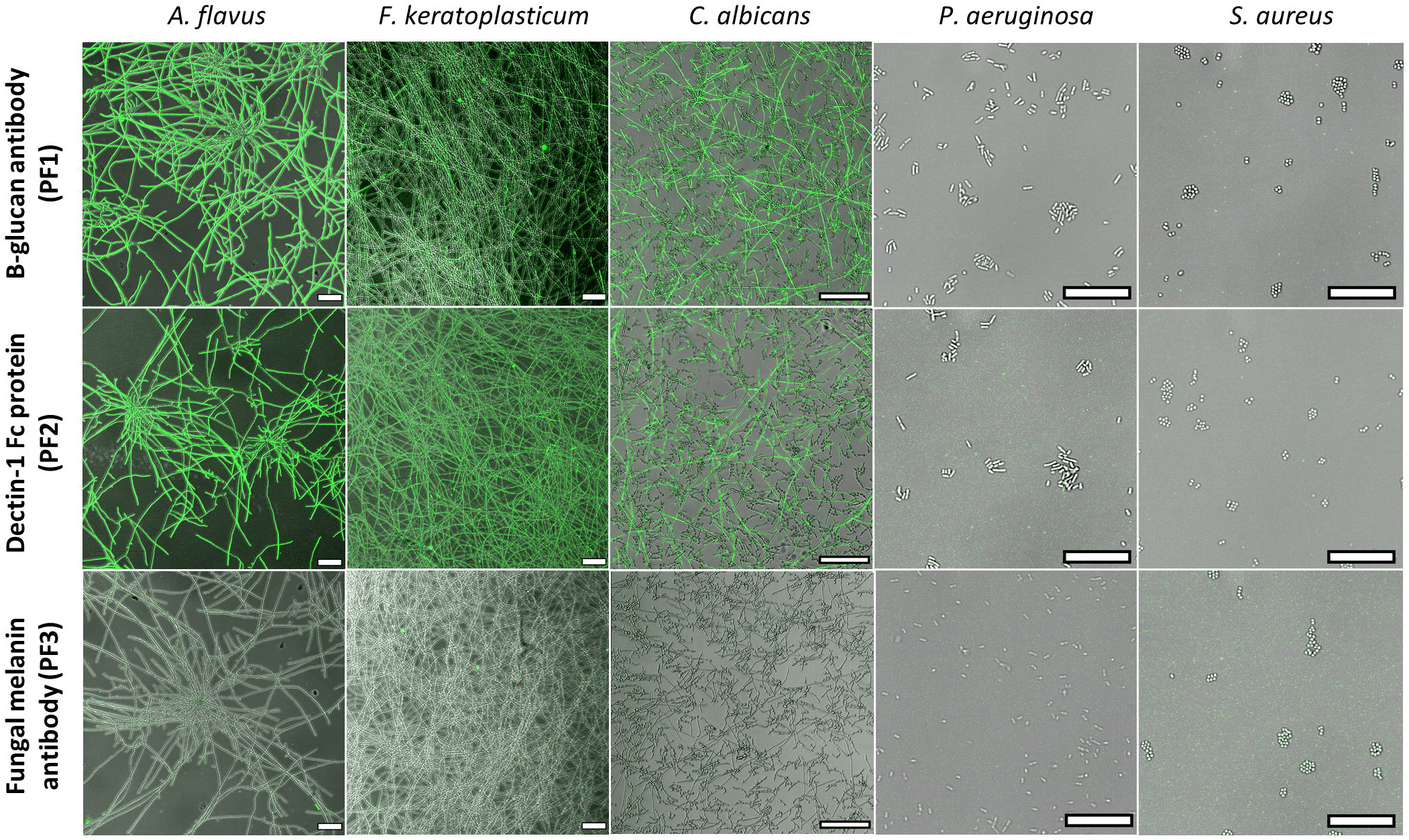
Screening for specific fungal binding capability of pan-fungal antibody candidates **(PF1-3)** by immunofluorescence. Representative images of immunofluorescence analysis are presented, showing a merged image of the respective antibody signal (green) and the bright field image (grey). Scale bar 100 µm fungi, *(A. flavus, F. keratoplasticum* and *C. albicans)*, 15 µm for bacteria *(P. aeruginosa* and *S. aureus)*.

### Semi-quantitative detection of zymosan and β-D-glucan standards with s-PF1-LFD and s-PF2-LFD

LFD strips with the two lead candidates were prepared in-house (**s-PF1-LFD** and **s-PF2-LFD**), and sensitivity was determined using positive control zymosan (a β-1,3-glucan derived from yeast). The linear detection range of **s-PF1-LFD** was between 10-75 µg/mL zymosan, after which a saturation of the signal was observed (**Figure 3a**). The **s-PF2-LFD** produced less intense signals, leading to a lower limit of detection at 50 µg/mL zymosan. The test bands were measured by visual inspection, and via a semi-quantitative approach measuring the ratio between the intensity of the test band and control band (T/C ratio), which has previously been shown to reduce ambiguity of visual inspection of LFDs.^19^ The T/C ratio trends were comparable to the visual scores with both lateral flow strip types. A significant positive correlation (*P*<0.0001) between zymosan concentration and signal intensity was observed for both in-house LFDs, when assessed by visual score and T/C ratio.

**Figure 3.**
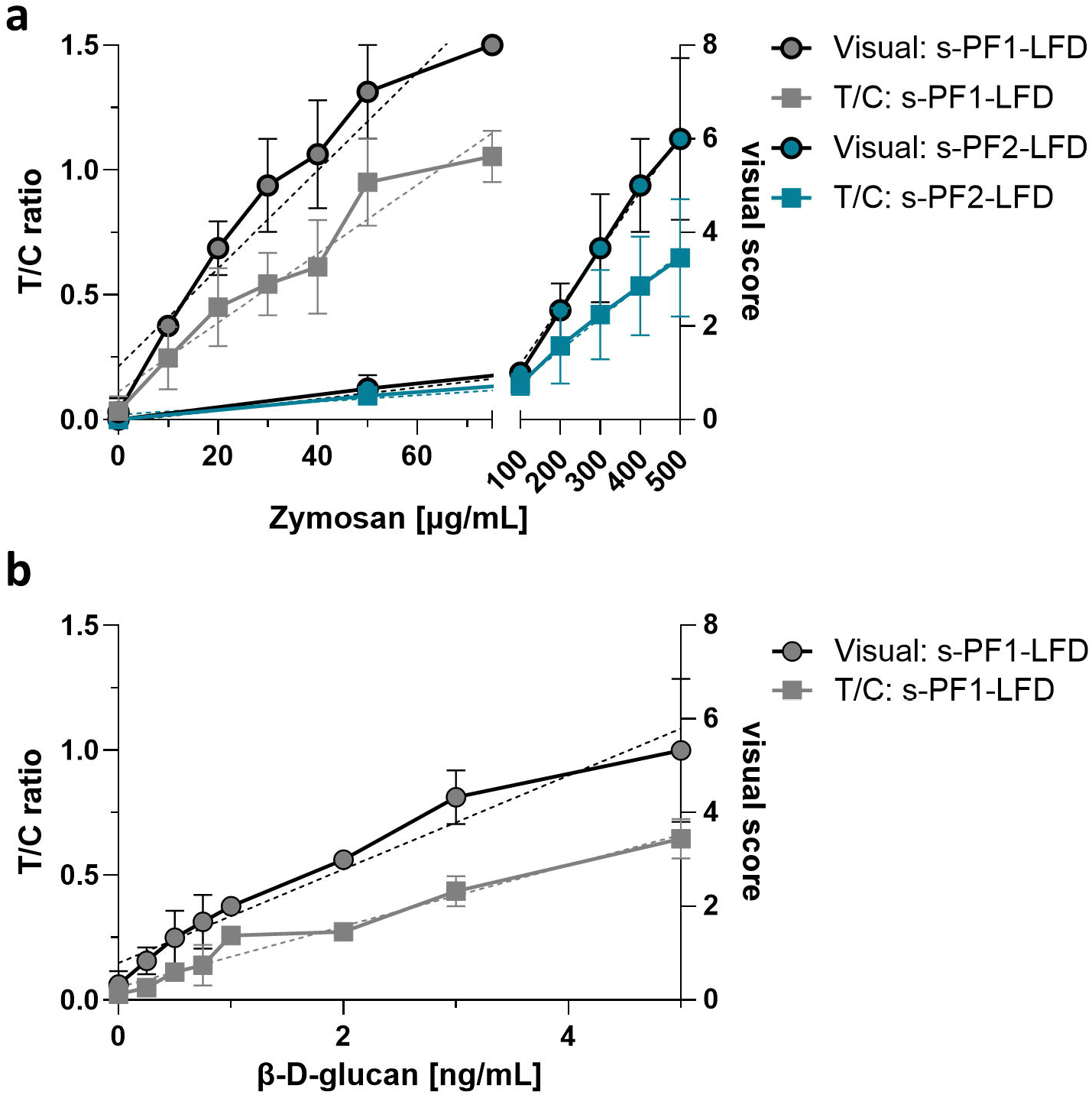
Visual and T/C ratio scoring of s-PF1-LFD and s-PF2-LFD strips evaluated with increasing concentrations of standards: **(a)** zymosan and **(b)** β-D-glucan (note: no signal was detectable for s-PF2-LFD with β-D-glucan). Mean values ± SD of three independent repeats depicted. Data analysed by simple linear regression (dashed lines), *P*<0.0001 for all conditions.

In contrast to zymosan, β-D-glucan was only detected with **s-PF1-LFD** (**Figure 3b**), with sensitivity in the low nanogram range (0.25-5 ng/mL). Again, the visual and measured T/C score trends were comparable, with a significant positive correlation (*P*<0.0001) between β-D-glucan concentration and signal intensity observed. Previous studies have reported β-D-glucan concentrations in the 1 ng/mL range from corneal scrapes of fungal keratitis patients,^25^ therefore the sensitivity of our test was within this clinically relevant range.

### s-PF1-LFD demonstrated superior sensitivity towards fungi compared to s-PF2-LFD

After determining the semi-quantitative range of detection using zymosan and β-D-glucan, we analysed fungal, bacterial, and buffer-only control samples with **s-PF1-LFD** and **s-PF2-LFD**. Buffer-only and bacteria samples assessed by the LFDs did not produce any positive test-bands, thus confirming specificity to the target fungi (**Supplementary Figure S1**). To measure LFD sensitivity, serial dilutions of fungal spores (*A. flavus* and *F. keratoplasticum*) and yeast (*C. albicans*) were prepared and grown into hyphae overnight. The hyphae were evaluated with **s-PF1-LFD** and **s-PF2-LFD**. The intensity of the resultant test bands was scored visually, independently by two scorers (**Figure 4a)** and by T/C ratio (**Figure 4b**). The T/C ratio and visual score of **s-PF1-LFD** increased in a concentration-dependent manner until saturating for samples seeded at 10^5^ spores or conidia mL^-1^. The lowest dilution of fungi detectable (seeded at 10^4^ spores mL^-1^) provided an equivalent of ∼1 ng/mL β-D-glucan (based on T/C ratio interpolated from **Figure 3b**) – in-line with the anticipated clinical range.^25^ In contrast **s-PF2-LFD** was two orders of magnitude less sensitive.

**Figure 4.**
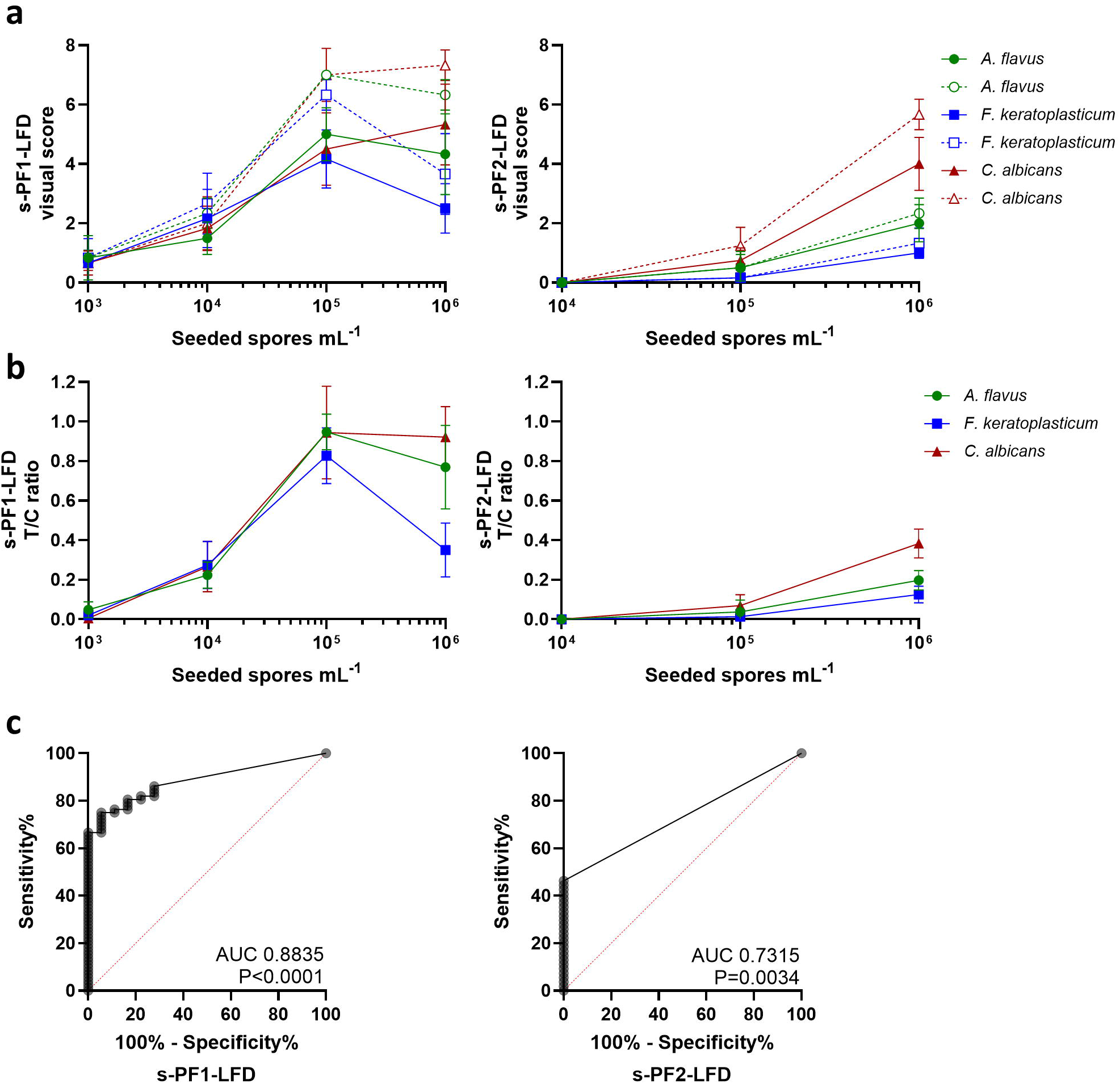
Fungal hyphae cultured from spores seeded at different concentrations were analysed by s-PF1-LFD and s-PF2-LFD. LFDs were **(a)** visually inspected by two independent, blinded scorers (closed symbols, solid line and open symbols, dashed line), and **(b)** T/C ratio. Both display mean SD values ± SD of three independent repeats. **(c)** Receiver operating characteristic (ROC) curves from the fungal measurements depicted in 3b (n=72 for s-PF1-LFD, n=54 for s-PF2-LFD), blanks and bacterial (*P. aeruginosa* and *S. aureus)* samples (n=18; 6 samples per blank and bacteria type).

Next, the overall sensitivity and specificity of **s-PF1-LFD** and **s-PF2-LFD** was determined by receiver operator characteristic (ROC) curves using the fungal (n=72) and non-fungal (bacteria and buffer (n=18)) T/C values (**Figure 4c**). Only **s-PF1-LFD** achieved an AUC of >0.8 (95% CI: 0.82-0.95, *P*<0.0001), indicating potential clinical utility.^26^ Based on the ROC curves, we selected a T/C cut-off of 0.112 to classify T/C ratios as “positive” or “negative”. **s-PF1-LFD** achieved a sensitivity of 0.75 (95% CI: 0.64-0.84), a specificity of 0.94 (95% CI: 0.74-0.99), and a likelihood ratio of 13.5. While **s-PF2-LFD** was highly specific (1.00, 95% CI: 0.82-1), sensitivity was low (0.46, 95% CI: 0.34-0.59).

### s-PF1-LFD was able to detect fungi present in samples collected from an *ex vivo* porcine cornea infection model

Due to the superior sensitivity of **s-PF1-LFD**, it was next evaluated for performance against samples collected in a clinically relevant manner, using an *ex vivo* model of microbial keratitis, with porcine cornea infected with fungi or bacteria. The samples were collected both by scrape (the clinical gold standard for obtaining corneal diagnostic specimens in tertiary clinics) and swab (which is minimally invasive, and would be suitable to be performed by an allied healthcare professional in multiple care settings).

The fungal samples obtained from the porcine cornea were positive when evaluated by **s-PF1-LFD**, resulting in strong bands when evaluated by visual inspection and T/C ratio (**Figure 5a**). In contrast, samples obtained from uninfected porcine corneas resulted in an average visual score of 1 or less, and extremely low T/C ratios, indicating no apparent test band.

**Figure 5.**
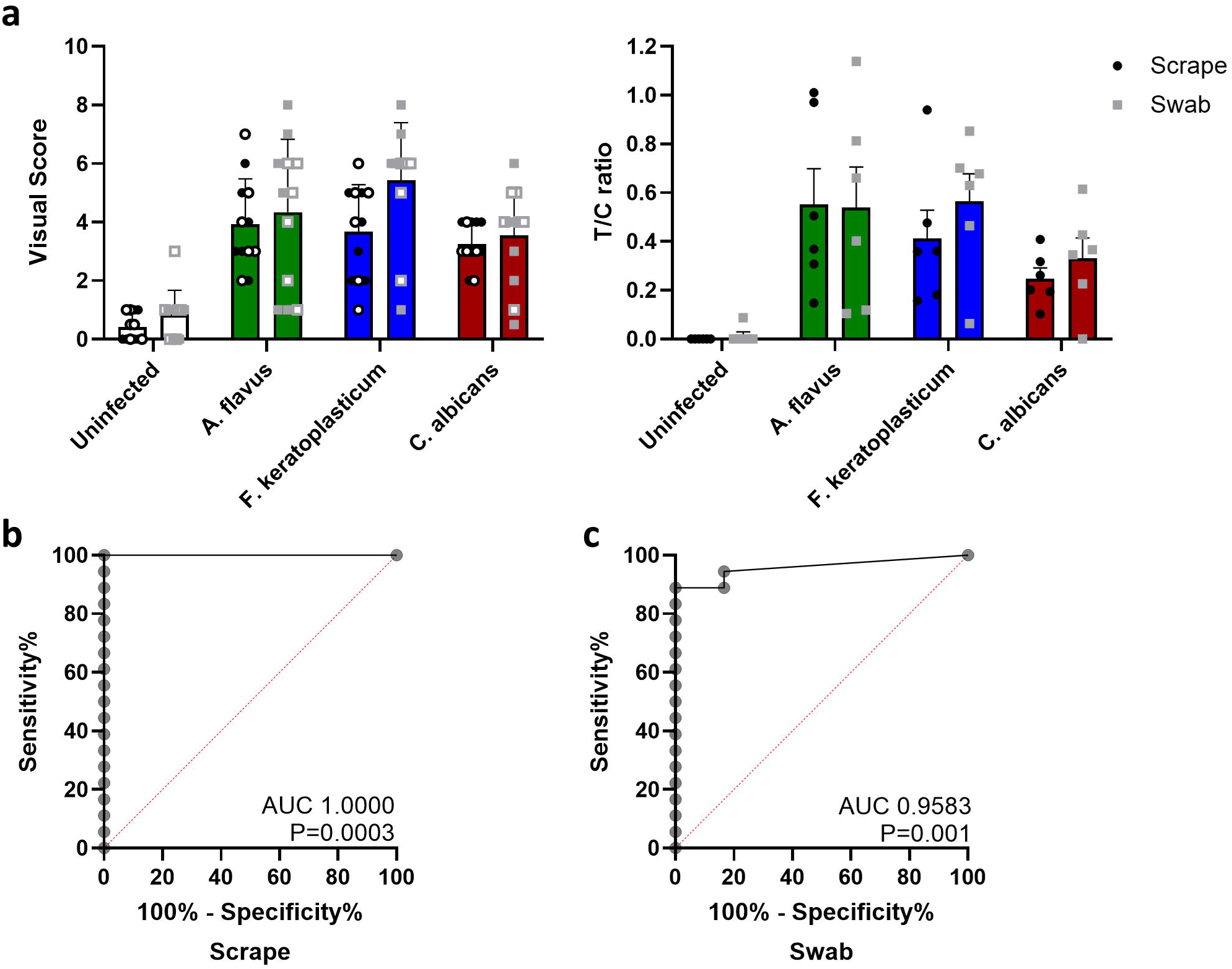
Scrapes (black dots) and swabs (grey squares) obtained from ex vivo porcine models of microbial keratitis were evaluated with s-PF1-LFD, and assessed by **(a)** visual scoring (with two independent, blinded scorers, open and closed symbols), and T/C band ratios. Samples collected over 3 independent experiments in duplicates. Error bars show mean ± SD. Receiver operating characteristic (ROC) curves obtained from **(b)** scrape and **(c)** swab T/C values of fungal keratitis (n=18) versus non-infected (n=6) samples.

The broad range in visual scores and T/C ratios for the samples obtained from the infected *ex vivo* porcine corneas is likely reflective of variation in levels of infection established in each cornea and/or the amount of pathogen that was retrieved during sample collection – further reflecting the clinical scenario. The pathogen load derived from scrapes and swabs from *F. keratoplasticum* or *A. flavus* infected eyes ranged from 10^1^-10^5^ CFU mL^-1^, whereas samples collected from *C. albicans* infected corneas were more consistent and ranged from 10^6^-10^8^ CFU mL^-1^. We found no significant difference between pathogen loads collected by either scrape or swab. From the scrape samples derived from the infected corneas, an AUC of 1.0 was obtained from the ROC curve analysis (**Figure 5b**), similar results were obtained from the swabs (AUC of 0.96, 95% CI: 0.88-1.0, **Figure 5c**). When applying a T/C cut-off value of >0.05, sensitivities and specificities of over 0.8 were achieved for both sample types (Table 1).

**Table 1:**
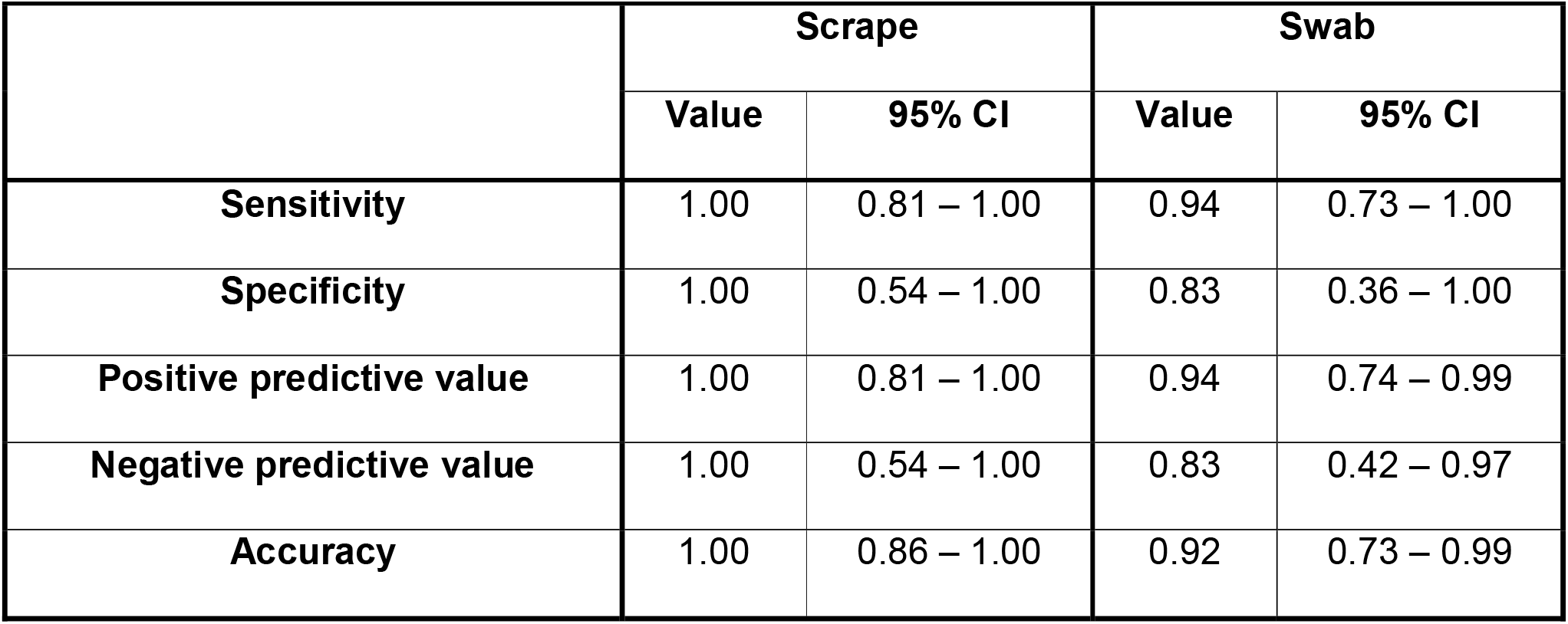
Diagnostic test evaluation of in-house lateral flow performance on scrape and swab samples.

### Evaluation of prototype p-PF1-LFD with zymosan and samples collected from an *ex vivo* porcine cornea infection model

In a next step of the development pipeline, we transferred our assay with **PF1** to a commercially-developed LFD system (producing **p-PF1-LFD**) to confirm compatibility with potential manufacture processes, and feasibility of translation to clinic. Here, we combined the LFD with a compatible commercial cube reader to enable real-time quantification of the test-line intensity, removing ambiguity of visual inspection and time-intensive steps of calculating T/C ratios.

In the first instance, **p-PF1-LFD** performance was confirmed with zymosan standards (**Figure 6a**). We found a comparable limit of detection and linear range for **p-PF1-LFD** that we did for our in-house **s-PF1-LFD**.

**Figure 6.**
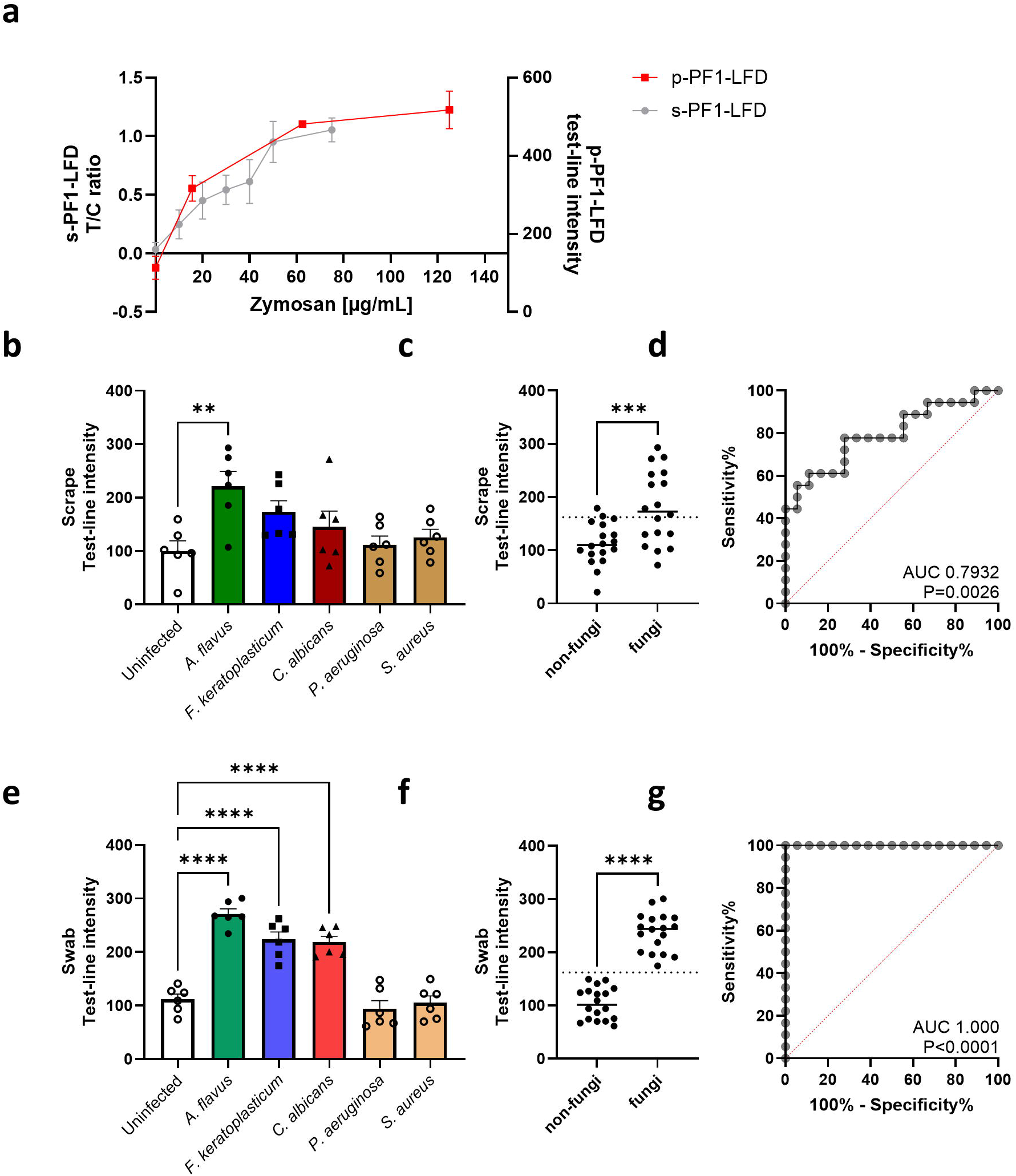
Lateral flow device prototypes **(p-PF1-LFD)** were evaluated using **(a)** zymosan, and **(b-d)** scrapes, and **(e-g)** swabs obtained from *ex vivo* porcine models of microbial keratitis. LFD signals were quantified using a cube reader after 20 minutes. Samples collected over 3 independent experiment in duplicates. Error bars show mean ± SD. **(b, e)** Ordinary one-way ANOVA, followed by Dunnett’s multiple comparisons test comparing to samples obtained from uninfected corneas. \*\**P*<0.01, \*\*\*\**P*<0.0001. **(c, f)** Cube-reader counts of all fungal (n=18) and non-fungal (n=18) cornea samples. Unpaired t-test. \*\*\**P*<0.001, \*\*\*\**P*<0.0001. **(d, g)** Receiver operating characteristic (ROC) curves obtained from **(c)** scrape and **(f)** swab test line intensities of fungal versus non-fungal cornea samples.

Next, the **p-PF1-LFD** prototype was evaluated using the *ex vivo* porcine cornea infection model, with samples collected by scrape (**Figure 6b-d**) and swab (**Figure 6e-g**). Overall, fungal samples (*A. flavus, F. keratoplasticum* and *C. albicans*) collected by minimally invasive swabs led to significantly higher test-band intensities compared to bacteria-infected and uninfected corneas. Samples collected by swab out-performed those collected by scraping when a test-band intensity threshold of 160 was applied, leading to excellent sensitivity and specificity for detecting a “fungal infection” irrespective of species (**Table 2**).

**Table. 2:** Diagnostic test evaluation of sandwich lateral flow performance on scrape and swab samples.

|  |  |  |  |  |
| --- | --- | --- | --- | --- |
|  | Scrape |  | Swab |  |
|  | Value | 95% CI | Value | 95% CI |
| **Sensitivity** | 0.56 | 0.31 – 0.78 | 1.00 | 0.81 – 1.00 |
| **Specificity** | 0.89 | 0.65 – 0.99 | 1.00 | 0.81 – 1.00 |
| **Positive predictive value** | 0.83 | 0.56 – 0.95 | 1.00 | 0.81 – 1.00 |
| **Negative predictive value** | 0.67 | 0.54 – 0.77 | 1.00 | 0.81 – 1.00 |
| **Accuracy** | 0.72 | 0.55 – 0.86 | 1.00 | 0.90 – 1.00 |

## Discussion

The current approach to diagnosing microbial keratitis (MK) by microbial culture and microscopy is not readily available in many places, and furthermore lacks the potential for easy application at primary care facilities where skilled laboratory personnel and laboratory resources are often unavailable. However, these facilities are often more widely available and accessible than tertiary care centres in regions with the highest burden of fungal keratitis.^27^ Consequently, there is an urgent need for simple, rapid and low-cost diagnostics for early differentiation of fungal MK cases from other causative pathogens for the rapid instigation of appropriate antimicrobials. Lateral flow device (LFD) technology is an increasingly appealing approach for development and deployment for disease diagnosis at primary care, however LFDs for MK are currently unavailable. We sought to address this technology gap by developing a pan-fungal LFD prototype, suitable for detection of key fungal species attributed to MK.

We evaluated the potential of antibodies targeting common fungal cell wall components for their sensitivity and specificity in an in-house LFD strip, as well as developing a preliminary commercial prototype with the lead antibody candidate. These were validated using pathogen samples from clinically relevant fungal and bacterial species, and further using samples collected by clinically relevant methods from *ex vivo* microbial keratitis models; we have previously demonstrated that antigen loads obtained from these models are comparable to human MK patient samples.^28^

Cell wall components shared between different fungal species, such as β-1,3-D-glucan, α-1,3-D-glucan, chitin, galactomannan or fungal melanin, are worth considering as target antigens for pan-fungal detection. However, chitin and α-1,3-glucan were excluded as no commercial antibodies were available at the time of our study. Nevertheless, in the future, non-commercial antibodies as developed in previous studies might facilitate LFD development with additional fungal antigen targets.^30,31^ In this study, galactomannan was excluded due to its relative specificity to *Aspergillus*^32^ and its potential for false-positivity caused by galactomannan contamination of some antibiotics.^33-35^ While a non-commercial fungal melanin antibody showed successful binding to multiple MK fungal isolates,^36^ this finding could not be confirmed with the anti-melanin antibody screened in our study. This could be due to differences in the fungal melanin antibody, or sample pre-treatment with cell-wall lysing enzymes in the study by Chongkae *et al*. Ultimately, a β-glucan antibody and a dectin-1 Fc conjugate remained for further screening in our in-house LFDs. Beta-D-glucan is a cell wall component of most pathogenic fungi, and readily used as the target antigen in existing diagnostic tests for invasive fungal infection, such as Fungitell® or β-glucan ELISAs. Its detection includes key pathogens of MK, such as *Aspergillus* spp., *Fusarium* spp. and *Candida* spp. It is noteworthy that species with low amounts of β-glucan antigen, such as *Mucorales, Cryptococcus neoformans* and *Blastomyces dermatitidis*, might be missed with a test based on β-glucan detection.^37^ These pathogens are however rarely involved in fungal keratitis.^38-40^

With our proof-of-concept, in-house developed LFD strips (**s-PF1-LFD**), we have demonstrated pan-fungal detection against target pathogens (*Aspergillus, Fusarium* and *Candida*) by detecting fungi but not bacteria grown in culture and from samples obtained from porcine cornea in a clinically applicable manner, achieving sensitivity and specificity of over 80%. Our approach further rules out any potential influence of mammalian host tissue towards the LFD readings. However, our in-house LFD strips require antibody conjugation steps, preparation of running buffer, further dilution steps of antibody conjugates, control and samples, and need to be stored at 4°C. Therefore, in a next step, the lead antibody (**PF1**) was used for prototype LFD generation. These LFDs are more applicable in point-of-care settings, as the sample simply needs to be obtained in the provided buffer and applied to the test. Prototype **p-PF1-LFD** achieved superior sensitivity and specificity with sample collected via the swabs compared to scrapes. Importantly, the sensitivity and specificity values of our pan-fungal LFDs are comparable to other diagnostic tests, such as smear microscopy (sensitivity of 0.61-0.94 and specificity of 0.91–0.97)^11^ or the Fungitell® test, which was recently evaluated for fungal MK diagnosis from tear samples (sensitivity of 0.96 and specificity of 0.83)^20^.

High LFD performance and compatibility with samples obtained from corneal swabs is essential. While scrape samples require skilled ophthalmologic personnel to obtain material from the corneal surface by scratching with a needle or blade, swab samples can be obtained from trained personnel at primary care centres, where diagnostic approaches are most urgently needed. We obtained similar results using both methods, which aligns with previous findings where MK scrape and swab samples performed with similar high sensitivity (0.94) when applied to an *Aspergillus* LFD.^19^ This confirms that swabs, as a minimally invasive sampling method, obtain sufficient antigen from MK patient eyes for LFD testing. The potential application of a pan-fungal LFD at the first healthcare interaction may enable initiation of appropriate antifungals at the earliest and most treatable stages of infection, improving patient outcomes.

While our data demonstrates potential utility of a pan-fungal LFD for the diagnosis of fungal keratitis, further optimisation and validation of the **p-PF1-LFD** prototype is required before study in a clinical environment. Despite the LFD prototype successfully differentiating fungal from non-fungal samples, we observed some level of non-specific binding at the test line with the uninfected and bacteria infected samples (∼100 counts). This creates ambiguity for reading the device by visual inspection, and may limit sensitivity in the field. Currently, the non-specific binding necessitates quantification of the test-band (via the cube reader) and a cut-off value to be applied to be able to differentiate fungal from non-fungal MK samples.

Furthermore, validation in a diagnostic study at a tertiary care facility is needed to allow for direct comparison of LFD results obtained from patient samples with gold-standard methods. Moreover, the application in primary care facilities needs to be evaluated to confirm ease of application at the point-of-care, and to verify whether patients at an earlier stage of fungal infection can be readily detected. Potential causes for false-positive results in a clinical setting, such as environmental contaminants, presence of β-glucan in cellulose used in surgical gauze,^41^ or contaminants in antibiotics ^42^ need to be examined. Environmental influences or contamination from make-up might be less severe than in other studies,^20^ as swab samples are examined instead of tears and hence more of the pathogen at the specific point of infection might be contained in the sample. Clinical studies would also evaluate the detectability of further MK fungal pathogens, as well as different strains, due to the broad spectrum of fungi presenting in clinics.

In conclusion, we have defined a promising lead antibody for the development of LFDs for application in MK diagnostic at the point-of-care setting, and further demonstrated preliminary feasibility. The key characteristics of these LFD include: (i) detecting *Candida, Fusarium* and *Aspergillus* fungal species; (ii) no cross-reactivity with common MK bacteria; (iii) detect antigen present in corneal scrapes and minimally invasive swabs; and (iv) provide a read-out within 20 minutes.

## Acknowledgements

This work was funded by a UKRI Future Leaders Fellowship (No. MR/V026097/1) and Sight Research UK (TRN003). Microscopy was carried out in the IRR Imaging facility at the University of Edinburgh. The authors acknowledge Browns Food Group (Quality Pork Processors, Ltd.) for the donation of ex vivo porcine eyes, and Lateral Dx Ltd. for their support in the development and manufacture of prototype lateral flow assays used in this study. The authors thank Dr Alex Kiang, Dr Florence Burté and Dr Kelvin Kah Wai Cheng for visual screening of LFD results. Figure 1 produced with BioRender (https://BioRender.com/ec988tr).

## References

1. Bartimote C, Foster J, Watson S. The spectrum of microbial keratitis: An updated review. The Open Ophthalmology Journal. 2019;13(1):100–130. doi:10.2174/1874364101913010100

2. Whitcher JP, Srinivasan M, Upadhyay MP. Corneal blindness: a global perspective. Bull World Health Organ. 2001;79(3):214–221.

3. Ung L, Acharya NR, Agarwal T, et al. Infectious corneal ulceration: a proposal for neglected tropical disease status. Bull World Health Organ. Dec 1 2019;97(12):854–856. doi:10.2471/BLT.19.232660

4. Lichtinger A, Yeung SN, Kim P, et al. Shifting trends in bacterial keratitis in Toronto: an 11-year review. Ophthalmology. Sep 2012;119(9):1785–1790. doi:10.1016/j.ophtha.2012.03.031

5. Shalchi Z, Gurbaxani A, Baker M, Nash J. Antibiotic resistance in microbial keratitis: ten-year experience of corneal scrapes in the United Kingdom. Ophthalmology. Nov 2011;118(11):2161–2165. doi:10.1016/j.ophtha.2011.04.021

6. Ung L, Bispo PJM, Shanbhag SS, Gilmore MS, Chodosh J. The persistent dilemma of microbial keratitis: Global burden, diagnosis, and antimicrobial resistance. Surv Ophthalmol. May-Jun 2019;64(3):255–271. doi:10.1016/j.survophthal.2018.12.003

7. Hoffman JJ, Burton MJ, Leck A. Mycotic Keratitis-A Global Threat from the Filamentous Fungi. J Fungi (Basel). Apr 3 2021;7(4)doi:10.3390/jof7040273

8. Brown L, Leck AK, Gichangi M, Burton MJ, Denning DW. The global incidence and diagnosis of fungal keratitis. Lancet Infect Dis. Mar 2021;21(3):e49–e57. doi:10.1016/S1473-3099(20)30448-5

9. Prajna NV, Srinivasan M, Lalitha P, et al. Differences in clinical outcomes in keratitis due to fungus and bacteria. JAMA Ophthalmol. Aug 2013;131(8):1088–1089. doi:10.1001/jamaophthalmol.2013.1612

10. Ting DSJ, Galal M, Kulkarni B, et al. Clinical Characteristics and Outcomes of Fungal Keratitis in the United Kingdom 2011-2020: A 10-Year Study. J Fungi (Basel). Nov 12 2021;7(11)doi:10.3390/jof7110966

11. Mills B, Radhakrishnan N, Karthikeyan Rajapandian SG, Rameshkumar G, Lalitha P, Prajna NV. The role of fungi in fungal keratitis. Exp Eye Res. Jan 2021;202:108372. doi:10.1016/j.exer.2020.108372

12. Moussa G, Hodson J, Gooch N, et al. Calculating the economic burden of presumed microbial keratitis admissions at a tertiary referral centre in the UK. Eye (Lond). Aug 2021;35(8):2146–2154. doi:10.1038/s41433-020-01333-9

13. Dalmon C, Porco TC, Lietman TM, et al. The clinical differentiation of bacterial and fungal keratitis: a photographic survey. Invest Ophthalmol Vis Sci. Apr 2 2012;53(4):1787–1791. doi:10.1167/iovs.11-8478

14. Wong TL, Ong ZZ, suresh l, et al. Expert Diagnostic Performance and Role of Corneal Sampling in Infectious Keratitis. Investigative Ophthalmology & Visual Science. 2025;66(8):602–602.

15. Sharma S, Kunimoto DY, Gopinathan U, Athmanathan S, Garg P, Rao GN. Evaluation of corneal scraping smear examination methods in the diagnosis of bacterial and fungal keratitis: a survey of eight years of laboratory experience. Cornea. Oct 2002;21(7):643–647. doi:10.1097/00003226-200210000-00002

16. Moledina M, Roberts HW, Mukherjee A, et al. Analysis of microbial keratitis incidence, isolates and in-vitro antimicrobial susceptibility in the East of England: a 6-year study. Eye (Lond). Sep 2023;37(13):2716–2722. doi:10.1038/s41433-023-02404-3

17. Thomas PA, Kaliamurthy J. Mycotic keratitis: epidemiology, diagnosis and management. Clin Microbiol Infect. Mar 2013;19(3):210–220. doi:10.1111/1469-0691.12126

18. Tuft SJ, Tullo AB. Fungal keratitis in the United Kingdom 2003-2005. Eye (Lond). Jun 2009;23(6):1308–1313. doi:10.1038/eye.2008.298

19. Gunasekaran R, Chandrasekaran A, Rajarathinam K, et al. Rapid Point-of-Care Identification of Aspergillus Species in Microbial Keratitis. JAMA Ophthalmol. Oct 1 2023;141(10):966–973. doi:10.1001/jamaophthalmol.2023.4214

20. Tawde Y, Das S, Singh S, et al. Evaluation of Fungitell (1,3)-β-D-glucan assay in tear samples for rapid diagnosis of fungal keratitis. Journal of Clinical Microbiology. 2024;62(12):e01200–01224. doi:doi:10.1128/jcm.01200-24

21. Koczula KM, Gallotta A. Lateral flow assays. Essays Biochem. Jun 30 2016;60(1):111–120. doi:10.1042/EBC20150012

22. Rousselot J, Millon L, Scherer E, et al. Detection of Mucorales antigen in bronchoalveolar lavage samples using a newly developed lateral-flow device. J Clin Microbiol. Jul 9 2025;63(7):e0022625. doi:10.1128/jcm.00226-25

23. Davies GE, Thornton CR. Development of a Monoclonal Antibody and a Serodiagnostic Lateral-Flow Device Specific to Rhizopus arrhizus (Syn. R. oryzae), the Principal Global Agent of Mucormycosis in Humans. J Fungi (Basel). Jul 21 2022;8(7)doi:10.3390/jof8070756

24. Grill FJ, Svarovsky S, Gonzalez-Moa M, et al. Development of a rapid lateral flow assay for detection of anti-coccidioidal antibodies. J Clin Microbiol. Sep 21 2023;61(9):e0063123. doi:10.1128/jcm.00631-23

25. Kaji Y, Hiraoka T, Oshika T. Increased level of (1,3)-beta-D-glucan in tear fluid of mycotic keratitis. Graefes Arch Clin Exp Ophthalmol. Jul 2009;247(7):989–992. doi:10.1007/s00417-008-1032-z

26. Çorbacioğlu Ş K, Aksel G. Receiver operating characteristic curve analysis in diagnostic accuracy studies: A guide to interpreting the area under the curve value. Turk J Emerg Med. Oct-Dec 2023;23(4):195–198. doi:10.4103/tjem.tjem_182_23

27. Rajendran VK, Raman R, Vardhan A, Gupta S, Bharath A, Ravilla TD. Impact of Vision Centres on achieving universal eye health coverage: prevalence, coverage and utilisation of eye care services in Southern India. British Journal of Ophthalmology. 2026;110(2):212–218.

28. Cheng KKW, Fingerhut L, Duncan S, Prajna NV, Rossi AG, Mills B. In Vitro and Ex Vivo Models of Microbial Keratitis: Present and Future. Prog Retin Eye Res. Jul 12 2024:101287. doi:10.1016/j.preteyeres.2024.101287

29. Yakushiji T, Inoue M, Koga T. Inter-serotype comparison of polysaccharides produced by extracellular enzymes from Streptococcus mutans. Carbohydr Res. Apr 15 1984;127(2):253–266. doi:10.1016/0008-6215(84)85360-4

30. Pisa D, Alonso R, Rabano A, Horst MN, Carrasco L. Fungal Enolase, beta-Tubulin, and Chitin Are Detected in Brain Tissue from Alzheimer’s Disease Patients. Front Microbiol. 2016;7:1772. doi:10.3389/fmicb.2016.01772

31. Figueiredo ABC, Fonseca FL, Kuczera D, Conte FP, Arissawa M, Rodrigues ML. Monoclonal Antibodies against Cell Wall Chitooligomers as Accessory Tools for the Control of Cryptococcosis. Antimicrob Agents Chemother. Nov 17 2021;65(12):e0118121. doi:10.1128/AAC.01181-21

32. Sherif R, Segal BH. Pulmonary aspergillosis: clinical presentation, diagnostic tests, management and complications. Curr Opin Pulm Med. May 2010;16(3):242–250. doi:10.1097/MCP.0b013e328337d6de

33. Walsh TJ, Shoham S, Petraitiene R, et al. Detection of galactomannan antigenemia in patients receiving piperacillin-tazobactam and correlations between in vitro, in vivo, and clinical properties of the drug-antigen interaction. J Clin Microbiol. Oct 2004;42(10):4744–4748. doi:10.1128/JCM.42.10.4744-4748.2004

34. Viscoli C, Machetti M, Cappellano P, et al. False-positive galactomannan platelia Aspergillus test results for patients receiving piperacillin-tazobactam. Clin Infect Dis. Mar 15 2004;38(6):913–916. doi:10.1086/382224

35. Aubry A, Porcher R, Bottero J, et al. Occurrence and kinetics of false-positive Aspergillus galactomannan test results following treatment with beta-lactam antibiotics in patients with hematological disorders. J Clin Microbiol. Feb 2006;44(2):389–394. doi:10.1128/JCM.44.2.389-394.2006

36. Chongkae S, Nosanchuk JD, Pruksaphon K, Laliam A, Pornsuwan S, Youngchim S. Production of melanin pigments in saprophytic fungi in vitro and during infection. J Basic Microbiol. Nov 2019;59(11):1092–1104. doi:10.1002/jobm.201900295

37. Dornbusch HJ, Groll A, Walsh TJ. Diagnosis of invasive fungal infections in immunocompromised children. Clin Microbiol Infect. Sep 2010;16(9):1328–1334. doi:10.1111/j.1469-0691.2010.03336.x

38. Khalili MR, Abtahi SMB, Atighehchian M, et al. Invasive Fungal Keratitis as an Uncommon Form of Mucormycosis Leading to Endophthalmitis: Report of Two Cases and Literature Review. Current Fungal Infection Reports. 2020/12/01 2020;14(4):384–390. doi:10.1007/s12281-020-00403-5

39. Tian J, Li D, Dai S, et al. In Vivo Confocal Microscopy Findings of a Rare Cryptococcus neoformans Keratitis. Ocul Immunol Inflamm. Dec 2024;32(10):2575–2578. doi:10.1080/09273948.2024.2386736

40. Lopez R, Mason JO, Parker JS, Pappas PG. Intraocular Blastomycosis: Case Report and Review. Clinical Infectious Diseases. 1994;18(5):805–807. doi:10.1093/clinids/18.5.805

41. Mennink-Kersten MA, Verweij PE. Non-culture-based diagnostics for opportunistic fungi. Infect Dis Clin North Am. Sep 2006;20(3):711-727, viii. doi:10.1016/j.idc.2006.06.009

42. Mennink-Kersten MASH, Warris A, Verweij PE. 1,3-β-D-Glucan in Patients Receiving Intravenous Amoxicillin–Clavulanic Acid. New England Journal of Medicine. 2006;354(26):2834–2835. doi:doi:10.1056/NEJMc053340

