## Supplemental Figure S1 for "Development and evaluation of a pan-fungal lateral flow device for the rapid identification of pathogen class in microbial keratitis"

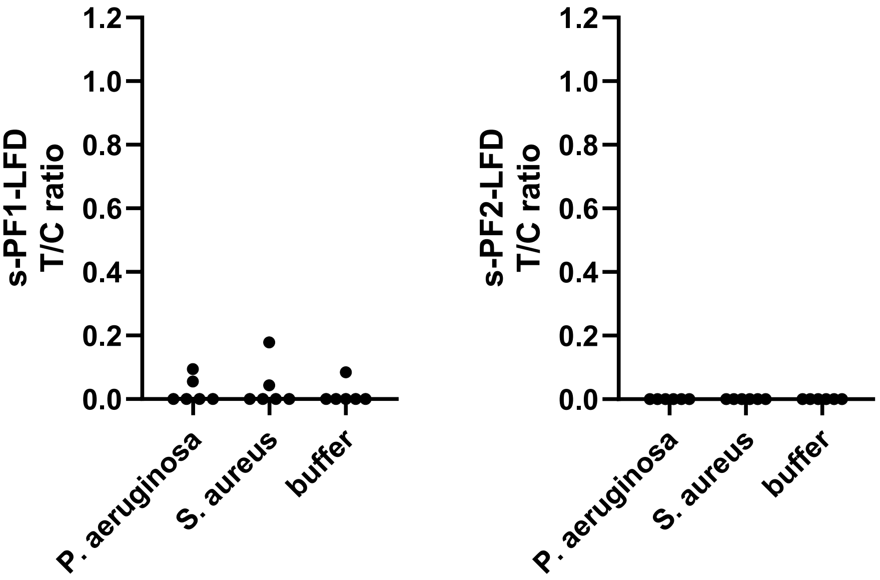


**Supplementary Figure S1: Assessment of s-PF1-LFD and s-PF2-LFD strips with cultured bacteria and buffer.** Signals obtained from *P. aeruginosa* (OD 0.1), *S. aureus* (OD 0.1), and running buffer were quantified using T/C ratio. All conditions display six values, of three independent repeats.
